# EvoPool: Evolution-Guided Pooling of Protein Language Model Embeddings

**DOI:** 10.64898/2026.02.02.703349

**Authors:** Navid NaderiAlizadeh, Rohit Singh

## Abstract

Protein language models (PLMs) encode amino acid sequences into residue-level embeddings that must be pooled into fixed-size representations for downstream protein-level prediction tasks. Although these embeddings implicitly reflect evolutionary constraints, existing pooling strategies operate on single sequences and do not explicitly leverage information from homologous sequences or multiple sequence alignments. We introduce EvoPool, a self-supervised pooling framework that integrates evolutionary information from homologs directly into aggregated PLM representations using optimal transport. Our method constructs a fixed-size evolutionary anchor from an arbitrary number of homologous sequences and uses sliced Wasserstein distances to derive query protein embeddings that are geometrically informed by homologous sequence embeddings. Experiments across multiple state-of-the-art PLM families on the ProteinGym benchmark show that EvoPool consistently outperforms standard pooling baselines for variant effect prediction, demonstrating that explicit evolutionary guidance substantially enhances the functional utility of PLM representations. Our implementation code is available at https://github.com/navid-naderi/EvoPool.

## 1. Introduction

Proteins are fundamental molecular machines of life, governing nearly all cellular processes through their structure, dynamics, and interactions. Understanding protein function from amino acid sequence alone is a long-standing challenge in biology.

In recent years, protein language models (PLMs) have emerged as a powerful paradigm for protein representation learning [1, 2]. Trained on massive corpora of protein sequences [3], PLMs leverage the abundance of evolutionary sequence data to learn rich contextual representations that capture biochemical and structural regularities. These models have demonstrated remarkable predictive performance across a wide range of downstream tasks, including protein structure prediction, functional annotation, and variant effect prediction [4–8].

Most PLMs operate in a sequence-only setting and produce residue-level (token-level) embeddings. However, many biologically relevant tasks, such as assessing protein function, stability, or fitness, are inherently defined at the protein level, rather than the residue level. This mismatch necessitates a pooling mechanism that aggregates token-level embeddings into a fixed-size, protein-level representation suitable for downstream prediction.

The simplest and most widely used pooling strategy is average pooling, which aggregates embeddings by taking their mean across residues. While effective and computationally convenient, average pooling treats all residues equally and ignores higher-order structures in the embedding space. Motivated by this limitation, several more sophisticated pooling mechanisms have been proposed, demonstrating improvements over average pooling across different settings [9–12].

Nevertheless, a critical limitation of existing pooling methods is that they operate on single sequences and cannot explicitly incorporate evolutionary information from homologous sequences or multiple sequence alignments (MSAs). Evolutionary relationships encoded in homologs are central to protein structure and function, and classical bioinformatics approaches have long relied on MSAs to identify conserved residues, functional motifs, and structural constraints [13–18]. While PLMs implicitly absorb some of this information during pretraining, they lack a mechanism to explicitly leverage homologous sequences at inference time. Although prior work has explored incorporating homologs at the PLM input level [19–24], explicitly integrating evolutionary information during the pooling stage has remained unexplored.

In this work, we introduce EvoPool, a self-supervised, constrained learning framework for evolution-guided pooling of PLM embeddings. EvoPool integrates homologous sequence information directly into the pooling process via an optimal transport formulation based on sliced Wasserstein distances. Our method constructs a fixed-size evolutionary anchor from an arbitrary number of homologous sequences and uses this anchor to aggregate query embeddings into an evolution-guided, protein-level representation. EvoPool is invariant to sequence length, the number of homologs, and homolog length, making it broadly applicable across diverse proteins and alignment depths.

We evaluate EvoPool on the ProteinGym benchmark [25], comprising approximately 2.4 million experimentally assayed substitution variants across 217 deep mutational scanning (DMS) experiments. Across three state-of-the-art PLM families, EvoPool consistently outperforms average pooling in zero-shot variant effect prediction, with gains that persist across biological function types, taxonomies, and MSA depth regimes. Moreover, we show that EvoPool provides complementary benefits even when applied on top of retrieval-augmented PLMs, demonstrating that homology-aware pooling captures evolutionary structure beyond what is captured at the model input.

## 2. Background and Related Work

### 2.1. Role of Homology in Protein Language Models

Most protein language models (PLMs) operate in a sequence-only setting, mapping a single amino acid sequence to a residue-level numerical representation. These models are typically trained on large sets of unaligned protein sequences using self-supervised objectives, thereby implicitly capturing evolutionary regularities in sequence data. Sequence-only PLMs such as ESM [5, 7, 26], ProtT5 [4], and ProGen [8, 27] have demonstrated strong performance across a wide range of downstream tasks, despite not explicitly incorporating information from homologous sequences during inference.

In parallel, several studies have explored the explicit incorporation of evolutionary information from homologous sequences to improve protein representations. A seminal contribution in this direction is the MSA Transformer [19], which operates directly on multiple sequence alignments (MSAs) and jointly models residue interactions across aligned homologs. By leveraging co-evolutionary patterns encoded in MSAs, MSA Transformer demonstrated substantial improvements in structure prediction and phylogenetic relationship identification [28]. However, its reliance on high-quality MSAs and quadratic scaling with alignment depth limits its applicability in large-scale or low-homology settings.

More recent models have explored alternative strategies for incorporating homology while mitigating these limitations. PoET [21, 22] introduces a retrieval-based framework that conditions sequence representations on a set of evolutionarily related sequences, sampled without requiring explicit alignment. This approach enables scalable incorporation of homologous information while maintaining flexibility in the number of retrieved sequences. Moreover, MSA Pairformer [23] presents a memory-efficient method for generating pairwise representations from MSA-based inputs, further improving structure prediction performance by efficiently modeling interactions both within and across homologous sequences.

E1 [24] represents a more recent retrieval-augmented PLM that unifies sequence-only and homology-aware inference within a single architecture. E1 retrieves a set of homologous sequences at inference time and conditions the encoding of the query sequence on this retrieved context, enabling adaptive use of evolutionary information depending on availability and computational budget. Importantly, E1 can operate in both sequence-only and retrieval-augmented modes, making it a flexible backbone for studying the effects of explicit homology integration.

Despite these advances, existing homology-aware PLMs primarily incorporate evolutionary information at the input stage. In contrast, relatively little attention has been paid to how homologous information should be incorporated during the pooling stage, where residue-level embeddings are aggregated into fixed-size protein-level representations. This distinction is particularly important for protein-level prediction tasks, such as variant effect prediction, where the choice of pooling mechanism plays a critical role in determining downstream performance. Our work complements prior efforts by explicitly focusing on evolution-guided pooling, providing a mechanism applicable to both sequence-only and homology-aware PLMs.

### 2.2. Pooling Protein Language Model Representations

To obtain fixed-size protein-level representations from residue-level embeddings produced by PLMs, most prior work has relied on simple aggregation operations, most notably average pooling [29–37]. Although average pooling is convenient to implement and computationally efficient, it assigns equal importance to all residues, whereas protein function and interactions are often governed by a small subset of critical residues. Moreover, for long proteins with hundreds of residues, average pooling can dilute informative signals, reducing discriminative power in the resulting representation.

These limitations have motivated the development of more sophisticated pooling mechanisms that aim to capture heterogeneity in residue importance and better preserve functional information. One such approach is light attention [9], which uses attention-based weighting to emphasize informative residues during aggregation. Similarly, BOM pooling [12] introduces a locality-aware pooling framework that combines average pooling and attention-based aggregation. Another example is PoolPaRTI [11], which leverages internal attention matrices from transformer layers and applies a PageRank-inspired procedure to estimate residue importance scores, thereby producing weighted protein-level embeddings that reflect structural and functional saliency. Most relevant to our work is PLM-SWE [10], which presents a principled pooling framework based on optimal transport. In this formulation, token-level embeddings and a set of learnable anchor elements are viewed as samples drawn from underlying probability measures, and pooling is achieved by computing sliced Wasserstein embeddings that align these distributions [38]. This approach provides a permutation-invariant, geometry-aware aggregation mechanism and has demonstrated consistent performance improvements across a range of downstream tasks, including drug-target interaction prediction and subcellular localization.

In this work, we build on the sliced Wasserstein embedding framework and extend it to explicitly incorporate homology information during pooling. We next provide a background on sliced Wasserstein embedding.

### 2.3. Sliced Wasserstein Embedding

The key idea behind sliced Wasserstein embedding (SWE) [38] is that it views token-level embeddings as samples drawn from an underlying high-dimensional probability measure. SWE generates constant-size permutation-invariant pooled embeddings for a set of representations, where the Euclidean distance between the pooled embeddings of two sets equals the sliced Wasserstein distance between the corresponding empirical measures in the original space.

Consider a set of query points 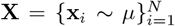, where each **x**_*i*_ ∈ ℝ^*d*^ denotes a *d*-dimensional token-level embedding, with *µ* denoting the underlying probability measure. Moreover, consider a set of *M* anchor embeddings 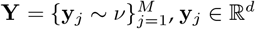, **y**_*j*_ ∈ ℝ^*d*^, drawn from an underlying anchor measure *ν*. We first map the query and anchor embeddings to a set of *L* 1-dimensional measures using a set of *L slices* 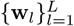, where each slicing direction sits on the unit hypersphere in ℝ^*d*^, denoted by 𝒮^*d*−1^.

For the *l*^th^ slice, SWE calculates the Monge coupling between the empirical sliced query and anchor distributions, i.e., 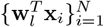 and 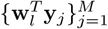, respectively. The *l*^th^ Monge coupling is a vector **z**_*l*_ = [*z*_*l*1_ … *z*_*lM*_]^*T*^ ∈ ℝ^*M*^ that contains the displacements from each sliced anchor element to its corresponding sorted sliced query element. For ease of notation, we assume that i) the query and anchor set have the same number of elements, i.e., *M* = *N*, and ii) the sliced anchor elements are already sorted in ascending order, i.e., 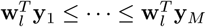. Then, the Monge coupling entries of the *l*^th^ slice are given by

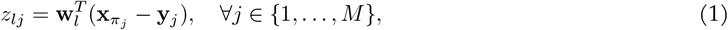

where *π* denotes the permutation indices for sorting the sliced query elements in ascending order, i.e., 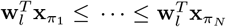. This operation generalizes, via additional sorting and interpolation, to cases in which the two assumptions above do not hold [38].

Collecting the Monge couplings across the *L* slices leads to the embedding **Z** ∈ ℝ^*M ×L*^. We use SWE to denote the end-to-end embedding pipeline, i.e., **Z** = SWE(**X**; **Y, W**), where **W** ∈ ℝ^*L*×*d*^ denotes the collection of the linear slicing directions. In cases where *M* is large, an optional weighting across the anchor elements, using a weight vector **u** ∈ ℝ^*M*^, can further reduce the final embedding size to an *L*-dimensional vector **z** = **Z**^*T*^ **u** ∈ ℝ^*L*^. The parameters of SWE, namely the anchor set **Y**, the slices **W**, and (optionally) the aggregation vector **u**, are trained using a downstream prediction loss.

While SWE has shown remarkable representation learning power in a variety of protein-related downstream tasks, such as protein-protein interaction [39] and subcellular localization [40], it is limited to supervised settings and does not provide a way for aggregating PLM embeddings in a task-agnostic manner [11]. Furthermore, it cannot account for evolutionary information. We build on the SWE framework to present a novel self-supervised pooling mechanism for PLM embeddings guided by homology information.

## 3. Proposed Method

Consider a query protein sequence ***s*** ∈ 𝒜^*N*^ with *N* residues, where 𝒜 denotes the amino acid alphabet. Moreover, consider an unordered set of *H* homologous sequences for ***s***, denoted by 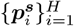. We pass both the query sequence ***s*** and its homologs through a sequence-only protein language model (PLM), which generates residue-level *d*-dimensional embeddings for the query and homologs, denoted by **S** ∈ ℝ^*N ×d*^ and 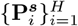, respectively, where the *i*^th^ homolog embedding, 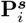, resides in 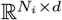, with *N*_*i*_ representing the length of the *i*^th^ homologous sequence. Note that we consider the general scenario in which i) the number of homologs, *H*, is arbitrary, and ii) each homolog may be of arbitrary length and does not need to have the same number of residues as the query sequence.

Our goal is to aggregate the residue-level query embedding **S** ∈ ℝ^*N ×d*^ into a fixed-size pooled embedding **z** ∈ ℝ^*d*^ that also incorporates the evolutionary information stemming from the homolog embeddings 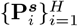.Formally, we seek a pooling function *f*_*θ*_, parameterized by a set of parameters *θ* ∈ Θ, that maps a variable-length query embedding and a variable-size collection of homologous sequence embeddings to a fixed-dimensional representation, i.e.,

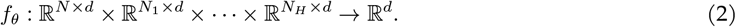

Importantly, this function should be invariant to the length of the query sequence, *N*, the number of the homologs *H*, and the lengths of the homologous sequences 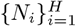.

Our key idea in this paper is to use the sliced Wasserstein embedding method described in §2.3 for the query sequence, *where the anchor is governed by the homolog embeddings*. We propose a two-level SWE operation in which the low-level SWE module aggregates homolog embeddings into a fixed-size anchor, which the high-level SWE module then uses to aggregate query embeddings into an evolution-guided, fixed-size representation of the query sequence.

Throughout this paper, we make use of *axis-aligned slices*, which imply that the number of slices equals the embedding dimensionality, i.e., *L* = *d*, and that the slices represent the standard basis vectors in ℝ^*d*^, i.e., **W** = **I**_*d*_, where **I**_*d*_ represents the *d* × *d* identity matrix. This choice removes the need to learn the slicer parameters, which we hypothesize better supports task-agnostic representation learning. The slicing directions may later be fine-tuned for downstream supervised tasks (see §5).

### 3.1. EvoPool: Evolutionary Aggregation of Protein Language Model Embeddings

#### Evolutionary Anchor Construction

We first use a low-level SWE module to map the homolog embeddings 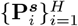 into a fixed-size *evolutionary anchor*. Using axis-aligned SWE, we use a learnable global anchor, **Y** ∈ ℝ^*M ×d*^, to pool each homolog embedding, i.e.,

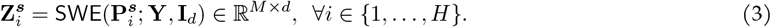

While the pooling operation in (3) maps each homolog into a fixed-size representation, we still need to summarize the *H* resulting representations into a fixed-size representation independent of the number of homologs. Homologs in multiple sequence alignments (MSAs) often exhibit substantial redundancy due to phylogenetic bias and database overrepresentation [41]. This implies that naively averaging pooled homolog embeddings can overemphasize densely sampled evolutionary lineages.

To address this issue, we use a *redundancy-aware weighting* scheme that operates directly on the pooled representations in (3). In particular, we estimate a soft local density around each embedding via

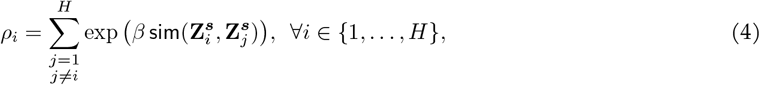

where sim(·,·) denotes cosine similarity between the *Md*-dimensional *flattened* embeddings and *β >* 0 is a hyperparameter that controls the sharpness of the density estimate. Each homolog is assigned an inverse-density weight *α*_*i*_ ∝ (*ρ*_*i*_ + 1)^−1^, normalized so that ∑_*i*_ *α*_*i*_ = 1. We then use the resulting weights to build our *evolutionary anchor* via weighted averaging of pooled homolog embeddings, i.e.,

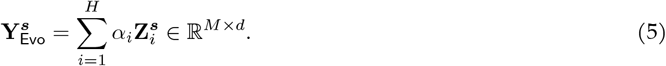

Intuitively, embeddings residing in dense clusters, corresponding to redundant or highly similar homologs, receive smaller weights, while isolated embeddings receive larger weights. This procedure serves as a soft, continuous analogue of classical redundancy correction in MSAs [42], ensuring that distinct evolutionary modes contribute approximately equally to the pooled representation. We detach the weights from gradient computation and treat them as frozen, allowing gradients to flow only through the pooled embeddings in (3).

#### Aggregating Query Embeddings

Given the evolutionary anchor 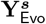 in (5), we then leverage it in another high-level SWE module to aggregate the query embeddings, i.e.,

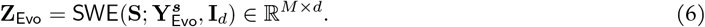

The aggregation in (6) is a bare-bones SWE pooling operation, which simply calculates the Monge couplings and does not learn any slicer or reference parameters. We finally use a weight vector *u* ∈ ℝ^*M*^ to derive the final embedding,

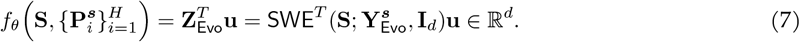

Figure 1 provides an overview of EvoPool. Note that the only learnable parameters in EvoPool include the global anchor for the low-level SWE and the weight vector in the high-level SWE, i.e., *θ* = (**Y, u**), comprising a total of *M* (*d* + 1) learnable parameters.

**Figure 1.**
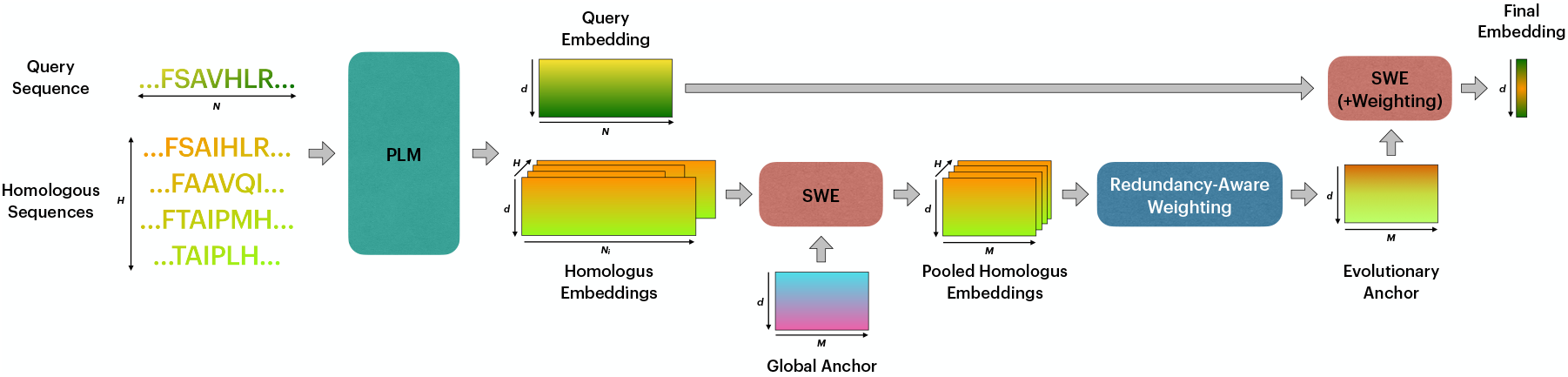
Overview of EvoPool. Given a query protein sequence and a variable number of homologous sequences, EvoPool aggregates residue-level embeddings generated by a protein language model (PLM) into a fixed-dimensional, evolution-guided protein representation. EvoPool first pools each homolog using a shared global anchor via sliced Wasserstein embedding (SWE), applies redundancy-aware weighting to construct an evolutionary anchor, and then aggregates the query embeddings against this anchor using a second SWE module. The resulting representation is invariant to query sequence length, number of homologs, and homolog sequence lengths, while explicitly incorporating evolutionary information during pooling.

### 3.2. Self-Supervised Training Procedure

We train EvoPool in a self-supervised manner. Our training goal is two-fold: (i) consistency of the query embedding given similar sets of homologs, and (ii) maximizing the diversity of embeddings within any collection of homologous sequences. We accomplish the former goal using a *contrastive self-supervised objective* [43, 44], while we encourage the latter goal using a *uniformity constraint* [45].

#### Contrastive Objective

Consider a batch of query sequence embeddings 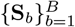. For each query **S**_*b*_, we randomly subsample two sets of homologs, whose embeddings we denote by 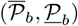, each possibly with a different depth. We then use the contrastive objective [44],

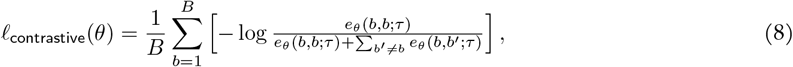

where *e*_*θ*_(*b, b*^*′*^; *τ*) is defined as

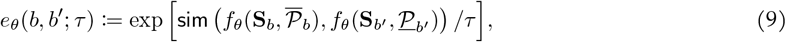

with *τ* denoting a temperature hyperparameter. Intuitively, minimizing the objective in (8) encourages the embeddings of a query sequence to stay unchanged given similar homolog embeddings, while simultaneously pushing the embeddings of different query sequences apart from each other in the embedding space.

#### Uniformity Constraints

To prevent representation collapse within sets of homologous sequences, we impose uniformity constraints on homologous embeddings. Consider two homologous embedding subsamples 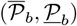 associated with a query sequence **S**_*b*_. Since embeddings in 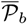 and 𝒫_*b*_ correspond to homologous sequences, each embedding in 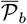 can itself be treated as a query and pooled against 𝒫_*b*_ using EvoPool (and vice versa). In particular, we can derive

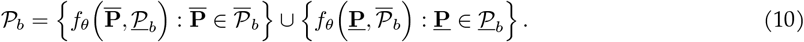

We then impose a uniformity constraint on the embeddings in 𝒫_*b b*_, i.e.,

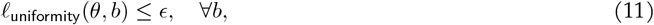

where *ϵ* denotes the uniformity bound hyperparameter, and the uniformity loss is defined as [45]

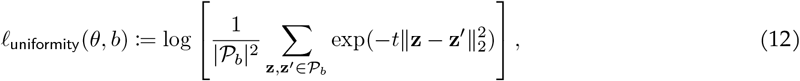

with *t* being a fixed hyperparameter that controls the range of the uniformity loss. The uniformity loss is always non-positive, and is approximately lower bounded by − 2*t*. Therefore, for active constraints, the upper bound *ϵ* should reside in the interval [ − 2*t*, 0). We set *t* = 2 in this paper as recommended in [45]. This constraint yields evolution-guided embeddings that encourage dispersion and prevent collapse across homologous sets.

#### Primal-Dual Self-Supervised Training

To solve the *constrained learning* problem that minimizes the contrastive objective in (8) subject to the uniformity constraints in (11), we leverage a primal-dual approach [46–48]. In particular, for each query sequence *b* in our training set, we introduce a non-negative dual multiplier *λ*_*b*_ ≥ 0, assembled in a vector ***λ***. Then, for a given batch of query embeddings 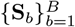, we form the Lagrangian

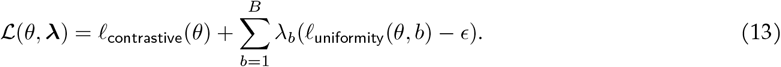

In the dual formulation, we need to minimize the Lagrangian over the primal model parameters *θ*, while maximizing it over the dual multipliers, i.e.,

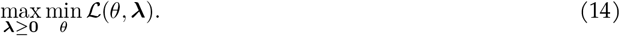

We solve the dual problem in (14) using iterative gradient descent-ascent. In particular, we update the model parameters via stochastic gradient descent, i.e.,

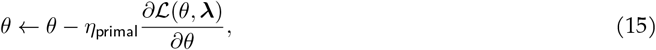

and we then update the dual multipliers via projected gradient ascent, i.e.,

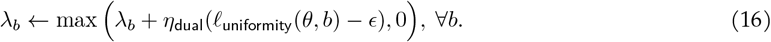

In (15)-(16), *η*_primal_ and *η*_dual_ denote the primal and dual learning rates, respectively. Note that the dual ascent updates in (16) cause the per-query dual multipliers to track the violations of the corresponding uniformity constraints. This implies that the dual variables increase dynamically for query samples whose homologous sequences undergo more severe internal representation collapse, whereas those for query samples whose homolog embeddings are sufficiently diverse remain close to zero.

## 4. Experiments

We train EvoPool on a set of 5000 randomly-selected MSAs from the UniClust30 clusters in OpenProteinSet [49]. We randomly subsampled homologs of size *H* ∈ {8, 9, …, 24} for each query sequence and set the uniformity constraint upper bound to *ϵ* = −*t* = −2. We use a batch size of 8 and train EvoPool for 40 epochs with a primal learning rate of 5 *×* 10^−4^ and a dual learning rate of 0.2. We set *β* = 50, *τ* = 0.1, and *M* = 128.

Once the self-supervised primal-dual training phase is complete, we evaluate EvoPool on the ProteinGym benchmark [25], a large-scale suite of deep mutational scanning (DMS) assays curated for rigorous assessment of protein variant effect predictors. We focus on ProteinGym’s 217 substitution assays, comprising approximately 2.4 million experimentally assayed variants derived from high-throughput functional screens.

We report results in the *zero-shot* variant effect prediction setting. In particular, to isolate and assess the quality of PLM representations, we score each variant using the *cosine similarity between the pooled wild-type and variant embeddings*, rather than likelihood-based approaches commonly employed for variant effect prediction [6, 50]. We measure performance using the Spearman correlation coefficient between the predicted scores and experimental fitness measurements. Because our evaluation is strictly zero-shot, we do not consider prior parameterized pooling methods that require task-specific training (e.g., [9, 10]). Instead, we adopt average pooling across residues as our primary baseline, as it is the most widely used and well-established pooling strategy in the PLM literature.

We consider three state-of-the-art backbones, namely ESM-2 650M [5], ESM-C 600M [26], and E1 600M [24]. While ESM-2 and ESM-C are sequence-only models, E1 allows both sequence-only and retrieval-augmented inference. Unless otherwise stated, we use the sequence-only inference mode for E1 to provide a fair comparison with ESM-2 and ESM-C. For each family, we use the available model scale that achieves the highest likelihood-based zero-shot performance on the ProteinGym DMS substitutions benchmark.

All our reported results are averages across two random seeds. Unless otherwise stated, we use *H* = 24 as the number of homologs during inference. We directly use the MSAs provided by ProteinGym to subsample homologs for all our variant effect prediction experiments.

### 4.1. Primal-Dual Training Results

We train EvoPool on top of the three PLM backbones using the primal-dual optimization procedure described in §3.2. Figure 2 summarizes the training dynamics, showing the evolution of the contrastive loss, the uniformity loss, and the empirical distribution of the learned non-zero dual multipliers at convergence.

**Figure 2.**
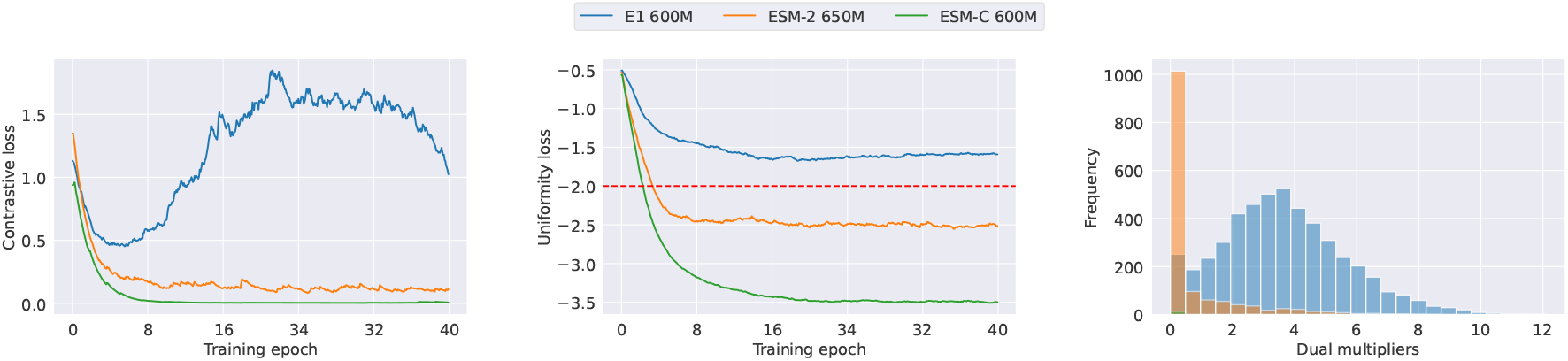
Convergence of (left) contrastive loss and (middle) uniformity loss, as well as (right) the distribution of final non-zero dual multipliers during the self-supervised training phase. The red dashed line in the middle plot represents the uniformity constraint threshold.

Across all models, the contrastive loss decreases rapidly in the early stages of training, indicating effective alignment between query representations across subsampled homologs and misalignment among distinct query representations. However, the final convergence behavior differs substantially across PLMs. ESM-C 600M achieves the lowest contrastive loss and converges smoothly, while ESM-2 650M follows closely with slightly higher residual loss. In contrast, E1 600M exhibits noticeably higher contrastive loss throughout training, suggesting a weaker ability to jointly optimize the two losses.

A similar pattern emerges for the uniformity constraint. As shown in the middle plot of Figure 2, ESM-C 600M consistently drives the uniformity loss well below the target threshold of *ϵ* = −2, indicating strong diversity of homologous embeddings and effective avoidance of representation collapse. ESM-2 650M also satisfies the constraint, though with a smaller margin. By contrast, E1 600M fails to reach the desired uniformity level, plateauing above the constraint boundary. This indicates that, for E1, enforcing uniformity conflicts more strongly with the contrastive objective.

These differences are further reflected in the learned dual multipliers. The distribution of dual variables for E1 600M is shifted toward significantly larger values, signaling sustained constraint violation at convergence. In comparison, both ESM-2 650M and ESM-C 600M exhibit substantially smaller dual multipliers, consistent with stable satisfaction of the imposed uniformity constraints.

### 4.2. EvoPool significantly outperforms average pooling for predicting the effect of substitutions

As shown in Table 1, EvoPool consistently improves zero-shot variant effect prediction performance across all PLM backbones, yielding substantial gains over average pooling and indicating that explicit evolutionary guidance can partially compensate for limited model capacity.

**Table 1:** Average zero-shot Spearman correlation coefficients (by function type) between experimental fitness values and predicted scores of EvoPool and average pooling (AvgPool) across three PLMs on the ProteinGym DMS substitution benchmark.

| Aggregation | PLM | Function Type |  |  |  |  |  |
| --- | --- | --- | --- | --- | --- | --- | --- |
|  |  | Average | Activity | Binding | Expression | Organismal Fitness | Stability |
| AvgPool | E1 600M | 0.0857 | -0.0267 | 0.0157 | 0.0065 | -0.0287 | 0.3277 |
|  | ESM-2 650M | 0.2124 | 0.1313 | 0.1244 | 0.1222 | 0.0999 | 0.4385 |
|  | ESM-C 600M | 0.3760 | 0.3581 | 0.2755 | 0.3433 | 0.3272 | 0.4735 |
| EvoPool | E1 600M | 0.2603 | 0.1789 | 0.1617 | 0.2429 | 0.1730 | 0.4394 |
|  | ESM-2 650M | 0.3423 | 0.2909 | 0.2345 | 0.2879 | 0.2714 | 0.4946 |
|  | ESM-C 600M | 0.4435 | 0.4371 | 0.3071 | 0.4226 | 0.3905 | 0.5421 |

When stratified by assay function type, EvoPool yields consistent improvements across all categories, with particularly pronounced gains for *activity, expression*, and *organismal fitness* assays. These tasks depend strongly on evolutionary constraints and functional context, suggesting that EvoPool’s homology-aware aggregation is especially effective when sequence-only embeddings are insufficient to capture functional variation. Our results also show improvements for binding and stability assays, indicating that EvoPool provides broad benefits across diverse biological readouts.

Table 2 further shows that EvoPool’s gains persist across organism types and MSA depth regimes. EvoPool substantially improves performance for human, eukaryotic, prokaryotic, and viral proteins. Moreover, EvoPool provides consistent improvements at low and medium MSA depths, where homologous information is sparse and standard pooling struggles, while still achieving strong performance at high MSA depths. This trend highlights EvoPool’s ability to effectively leverage limited evolutionary context without relying on deep alignments.

**Table 2:**
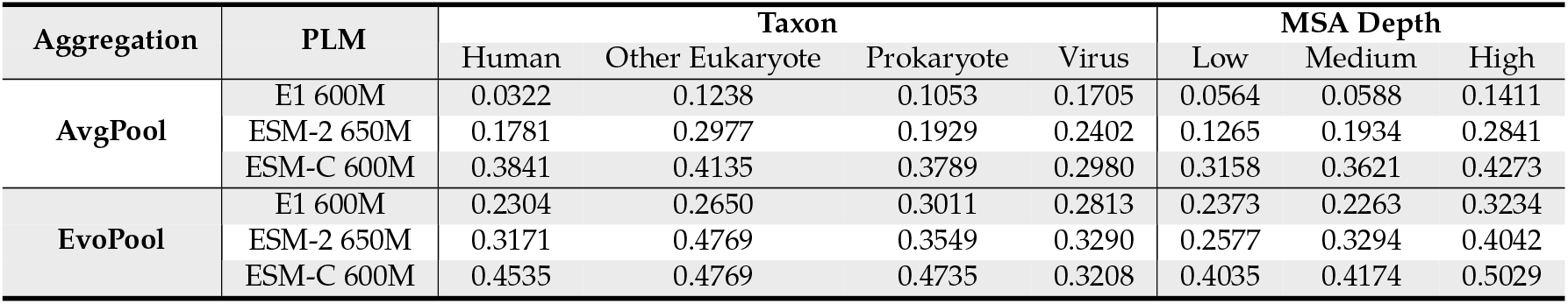
Average zero-shot Spearman correlation coefficients (by taxonomic group and MSA depth) between experimental fitness values and predicted scores of EvoPool and average pooling (AvgPool) across three PLMs on the ProteinGym DMS substitution benchmark.

| Aggregation | PLM | Taxon |  |  |  | MSA Depth |  |  |
| --- | --- | --- | --- | --- | --- | --- | --- | --- |
|  |  | Human | Other Eukaryote | Prokaryote | Virus | Low | Medium | High |
| AvgPool | E1 600M | 0.0322 | 0.1238 | 0.1053 | 0.1705 | 0.0564 | 0.0588 | 0.1411 |
|  | ESM-2 650M | 0.1781 | 0.2977 | 0.1929 | 0.2402 | 0.1265 | 0.1934 | 0.2841 |
|  | ESM-C 600M | 0.3841 | 0.4135 | 0.3789 | 0.2980 | 0.3158 | 0.3621 | 0.4273 |
| EvoPool | E1 600M | 0.2304 | 0.2650 | 0.3011 | 0.2813 | 0.2373 | 0.2263 | 0.3234 |
|  | ESM-2 650M | 0.3171 | 0.4769 | 0.3549 | 0.3290 | 0.2577 | 0.3294 | 0.4042 |
|  | ESM-C 600M | 0.4535 | 0.4769 | 0.4735 | 0.3208 | 0.4035 | 0.4174 | 0.5029 |

### 4.3. EvoPool’s gains increase with sequence length, except for ESM-C

We further analyze how EvoPool’s performance gains over average pooling vary with protein sequence length. Figure 3 plots the absolute Spearman correlation improvement of EvoPool relative to average pooling as a function of sequence length for each PLM backbone across assays with at most 1000 residues.

**Figure 3.**
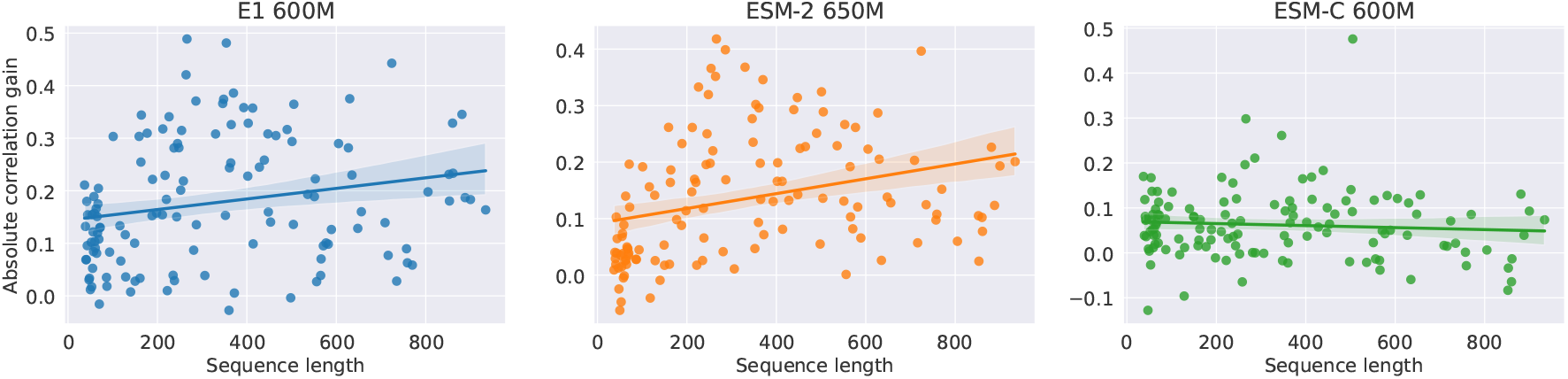
Absolute Spearman correlation gains of EvoPool compared to average pooling as a function of sequence length across three PLMs on the ProteinGym DMS substitution benchmark.

For E1 600M and ESM-2 650M, EvoPool’s gains exhibit a clear positive trend with increasing sequence length. In particular, longer proteins tend to benefit more from homology-aware pooling, suggesting that explicit evolutionary aggregation becomes increasingly important as sequences grow longer and functional dependencies span larger regions of the protein. This behavior is consistent with the limitations of simple average pooling, which dilutes informative residue-level signals in long sequences.

In contrast, EvoPool’s gains for ESM-C 600M remain stable across sequence lengths. This flatter trend reflects ESM-C’s potentially stronger ability to capture long-range dependencies directly within its residue-level representations, reducing, but not eliminating, the benefit of evolutionary pooling for longer proteins. EvoPool still provides consistent improvements across the full length range, indicating that homology-aware aggregation complements even PLMs with enhanced long-context modeling capabilities.

### 4.4. Leveraging homology both at input retrieval and during pooling improves zero-shot prediction

To isolate the contributions of homology information at different stages, we conduct additional experiments using E1, a retrieval-augmented PLM that incorporates homologous sequences at the input level. Due to the computational cost of retrieval-augmented inference, we restrict this analysis to 46 ProteinGym DMS substitution assays with sequence lengths of at most 100 amino acids and no more than 2000 variants per assay. We also limit the number of sampled homologs to 8 per query sequence.

Figure 4 compares zero-shot variant effect prediction performance when homologous sequences are used only during input retrieval, followed by average pooling, versus when they are additionally incorporated during pooling via EvoPool. Quite interestingly, across most assays, incorporating homology at both stages yields consistent, often substantial improvements over retrieval alone. Notably, the gains from EvoPool persist even when retrieval-augmented representations are available, indicating that pooling-time aggregation captures complementary evolutionary structure that is not fully exploited at the input level. This effect is particularly pronounced for assays where retrieval-only performance is modest, suggesting that EvoPool can amplify weak evolutionary signals by enforcing structured aggregation over homologous embeddings. We note that, for E1, EvoPool was trained using sequence-only embeddings for both queries and homologs during the primal-dual optimization phase. We therefore expect that training or fine-tuning EvoPool directly on retrieval-augmented embeddings could further enhance its gains in E1’s retrieval-augmented inference mode.

**Figure 4.**
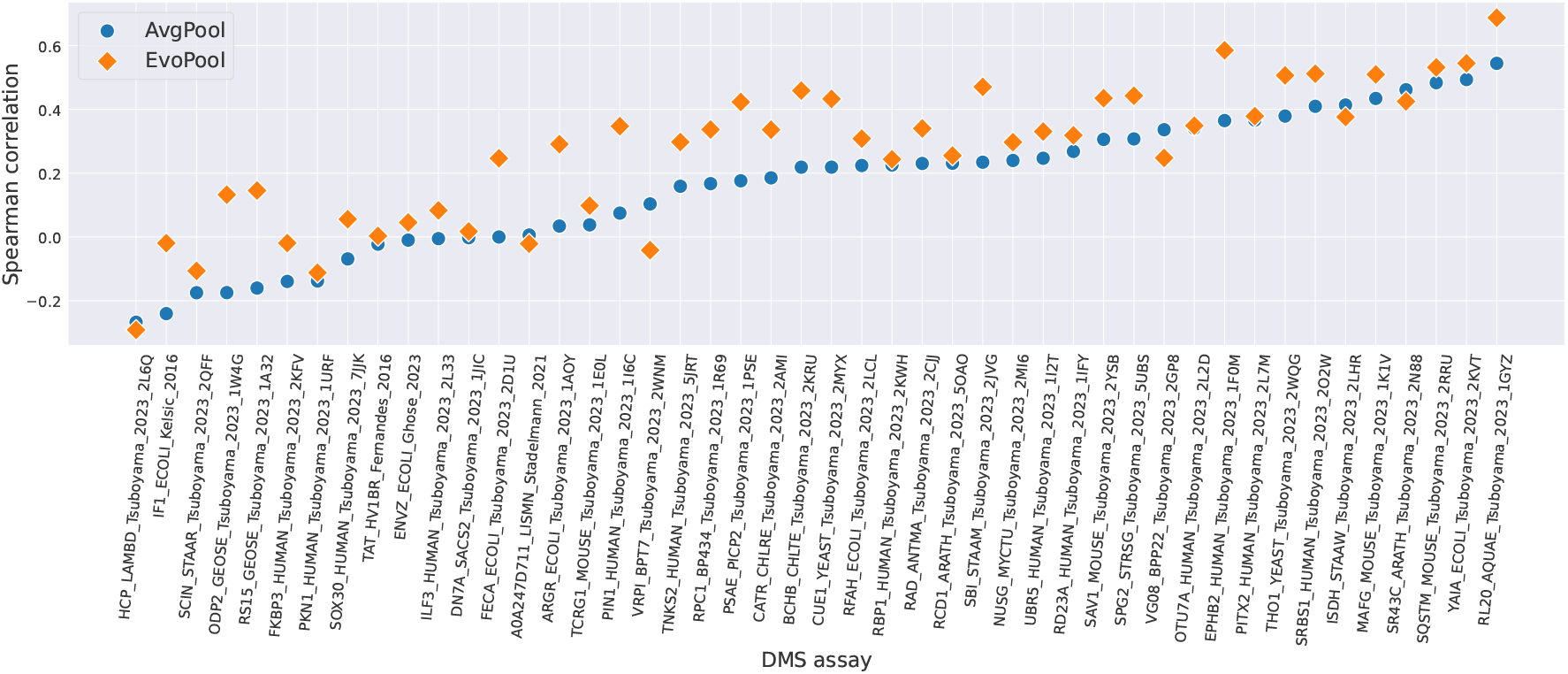
Comparison of EvoPool and average pooling (AvgPool) for the E1 600M PLM in the retrieval-augmented mode with 8 homologs on 46 ProteinGym DMS assays with at most 100 residues and 2000 variants per assay.

These results demonstrate that homology-aware pooling provides benefits beyond retrieval augmentation and that explicitly integrating evolutionary information during pooling remains valuable even for PLMs designed to incorporate homologs at inference time.

## 5. Discussion

We introduced EvoPool, a self-supervised, optimal-transport-based pooling framework that explicitly integrates evolutionary information from homologous sequences into protein language model (PLM) representations. EvoPool addresses a fundamental limitation of standard pooling strategies, which operate on single sequences and cannot exploit homology signals that are central to protein function. By constructing a fixed-size evolutionary anchor from an arbitrary number of homologs and using sliced Wasserstein aggregation to pool query embeddings, EvoPool provides a principled mechanism for generating evolution-guided protein representations.

Extensive evaluations on the ProteinGym DMS substitution benchmark show that EvoPool consistently outperforms average pooling across multiple PLM families, biological function types, organismal groups, and MSA depth regimes. Our results illustrate that EvoPool is particularly effective in challenging settings, including proteins with limited homologous information and long sequences where naive pooling dilutes informative residue-level signals. Additional experiments with retrieval-augmented inference further demonstrate that incorporating evolutionary information during pooling yields benefits beyond input-level retrieval.

Several extensions of EvoPool offer promising directions for future research. In this work, we relied on random subsampling of homologs from MSAs for both training and inference. More informed sampling strategies, such as diversity-aware or PoET-style evolutionary sampling [21, 22, 24], could further improve the quality of the evolutionary anchor while reducing computational cost. In addition, while our empirical study focused on substitution DMS assays, extending the evaluation to indel benchmarks would provide a more comprehensive assessment of EvoPool’s ability to capture evolutionary constraints involving larger structural perturbations.

From a methodological perspective, we adopted axis-aligned slicing directions for the sliced Wasserstein embedding, yielding a task-agnostic, parameter-efficient pooling mechanism. While this choice simplifies training and improves robustness, learning or adapting slicing directions to specific downstream tasks may further enhance performance in supervised or semi-supervised settings.

